# Intratumoral B7H3:CD3 Bispecific T-cell Engager Drives Localized T-cell Accumulation in Canine Sarcoma Patients

**DOI:** 10.64898/2026.05.27.728355

**Authors:** Yusuke Suita, Lisa S. Ang, Kenneth Brasel, Shelli M. Morris, Emily J. Girard, Alison M. Williams, Steven C. Chen, Ian Blumenthal, Natasha M. Hottmann, Jennifer Heusser, Andrew J. Mhyre, Cole A. DeForest, Peter Moore, Jason P. Price, Janean Fidel, James M. Olson

**Author notes:** Co-2nd authors.

## Abstract

**Background:** Bispecific T-cell Engagers (TCEs) targeting B7H3 (CD276) show promise for solid tumors but are limited by systemic toxicities and poor tumor penetration. Intratumoral (IT) delivery is proposed as a solution, but the safety and spatial pharmacodynamics (PD) remain poorly defined in these malignancies. Spontaneous canine tumors serve as a highly translatable model for human therapeutic development due to its clinical, genetic, and immunological similarities to human patients. This study evaluates the feasibility of an IT-delivered B7H3:CD3 TCE in a trial that enrolls companion dogs with solid tumors.

**Methods:** We engineered a canine B7H3:CD3 TCE and validated its ability to induce T-cell activation and T-cell mediated cytotoxicity *in vitro* on several B7H3-expressing canine tumor cell lines. Two STS canine patients received intratumoral columnar injections of the TCE and saline (internal control) at fixed distance of 1.5cm using a custom-engineered multi-needle assembly. Safety was evaluated by physical examinations and hematological and biochemical changes in peripheral blood. PD response was analyzed by H&E and immunohistochemistry.

**Results:** *In vitro* assays validated the cytotoxicity of the B7H3:CD3 TCE on B7H3^+^ canine tumor cell lines. TCE IT administration (7.83 μg / 148.2 pmol) was well tolerated with no adverse events greater than Grade 1 and no evidence of systemic cytokine release or organ toxicity. Immunohistochemistry of tumors collected 7 days after TCE administration revealed a significant five-fold increase in CD3^+^ T-cell density at the TCE injection site (within 0.5 cm radius) compared to internal saline controls.

**Conclusions:** This study demonstrated the feasibility of evaluating pharmacodynamic response to IT delivery of B7H3:CD3 TCE, namely local T-cell accumulation. T-cell localization around the TCE injection site supports our hypothesis that effective IT immunotherapy might require enhanced volumetric coverage using multi-needle injections and/or co-stimulatory strategies to convert T-cell localization into a robust, sustained anti-tumor response.

## Introduction

B7H3 (CD276), a member of the B7 family of immunomodulatory ligands, has emerged as a compelling target for cancer immunotherapy due to its overexpression in pediatric and adult solid tumors, including soft tissue sarcomas (STSs) (1, 2). In humans and dogs, high B7H3 expression correlates with advanced tumor grade, metastatic potential, and poor survival (3, 4). However, the clinical translation of B7H3-targeted immunotherapy is frequently hampered by a narrow therapeutic window (5).

While B7H3 is tumor-enriched, low-level expression in healthy peripheral tissues (e.g., liver) poses a risk for “on-target/off-tumor” toxicities (6). In a Phase I/II trial, intravenous injection of a B7H3-targeted antibody (Enoblituzumab) plus PD-1 inhibitor therapy for advanced solid tumors caused treatment-related adverse events in 87.2% of patients, with 28.6% experiencing Grade 3 or higher toxicities (7). Systemic administration of CD3-engaging TCEs often trigger cytokine release syndrome (CRS), a systemic inflammatory response characterized by fever, hypotension, and multi-organ dysfunction (8). These safety concerns necessitate dose de-escalations that often render therapy sub-therapeutic within the immunosuppressive tumor microenvironment (TME). Furthermore, rapid systemic clearance and poor extravasation into bulky, high-pressure sarcoma masses limit therapy exposure (9). Thus, there is an urgent clinical need for B7H3:CD3 TCE delivery strategies that achieve high local T-cell densities without triggering systemic toxicity. Intratumoral (IT) delivery has been shown to maximize tumor exposure and minimize on-target/off-tumor exposure (10–12). By bypassing systemic pharmacokinetics and biodistribution, direct injection ensures that therapeutic concentrations of TCEs are achieved locally and limits exposure to healthy tissues to the TCE, thereby reducing the potential for systemic toxicity. However, for IT delivery to be effective, a sufficient density of activated cytotoxic T cells must distribute throughout the tumor volume. In bulky solid tumors, high interstitial fluid pressure, a dense, desmoplastic stroma, and vascular dysregulation create significant physical barriers to therapy diffusion and T-cell cellular motility (13, 14). Without understanding the spatial pharmacodynamics (PD) of a single-injection T-cell engager (TCE), IT delivery risks insufficient saturation or localized T cell overloading that may trigger unintended toxicity or T cell dysfunction.

Spontaneous canine malignancies represent a high-fidelity translational model for evaluating novel human immunotherapies (15). Canine tumors arise spontaneously in immune-competent hosts and share similar genomic instabilities, histopathological complexity, and intratumoral heterogeneity found in human patients. Pet dogs are exposed to diverse environmental antigens, which result in a mature and educated immune system that more accurately reflects human T-cell dynamics rather than the controlled, pathogen-free environments of laboratory mice. Crucially, the human-comparable physical scale allows for the mapping of a therapy’s spatial distribution. Given the inherent inter-patient heterogeneity in T-cell density and baseline activation, we used an internally controlled design to isolate the effects of the TCE. By using a spatially fixed multi-needle assembly to deliver a localized dose of B7H3:CD3 TCE, we used each patient as their own internal control to determine whether T-cell enrichment is a global or localized phenomenon in the tumor (16, 17).

This study demonstrated that a localized B7H3:CD3 TCE dose achieves a significant five-fold increase in T cells within a 1.5cm radius from the TCE injection site compared to a saline injection site one week post-injection. The injection was well tolerated with no systemic toxicities or acute injection site reactions observed. With these findings supporting the feasibility of B7H3:CD3 TCE IT delivery, a clinical trial has been launched to assess B7H3:CD3 TCE plus co-stimulatory TCEs in a dose escalation design to maximize T-cell activation and anti-tumor activity at tolerated doses.

## Results

### *In Vitro* Potency and Antigen-Specific Cytotoxicity of B7H3:CD3 TCE

To address the feasibility of conducting a local immunotherapy trial in canine cancer patients, we first engineered a TCE comprised of single-chain variable fragments (scFv) that recognize canine B7H3 and CD3 and that are connected by a flexible polypeptide linker (Fig. 1A). This compact, single-chain format was selected for its dose and target-dependent *in vitro* activity and reduced molecular weight, which facilitates superior tissue penetration compared to full-length IgG formats (18). Furthermore, the smaller molecular size provides a favorable safety profile; in the event of systemic circulation, the rapid renal clearance minimizes prolonged systemic exposure and the associated CRS risk.

**Figure 1.**
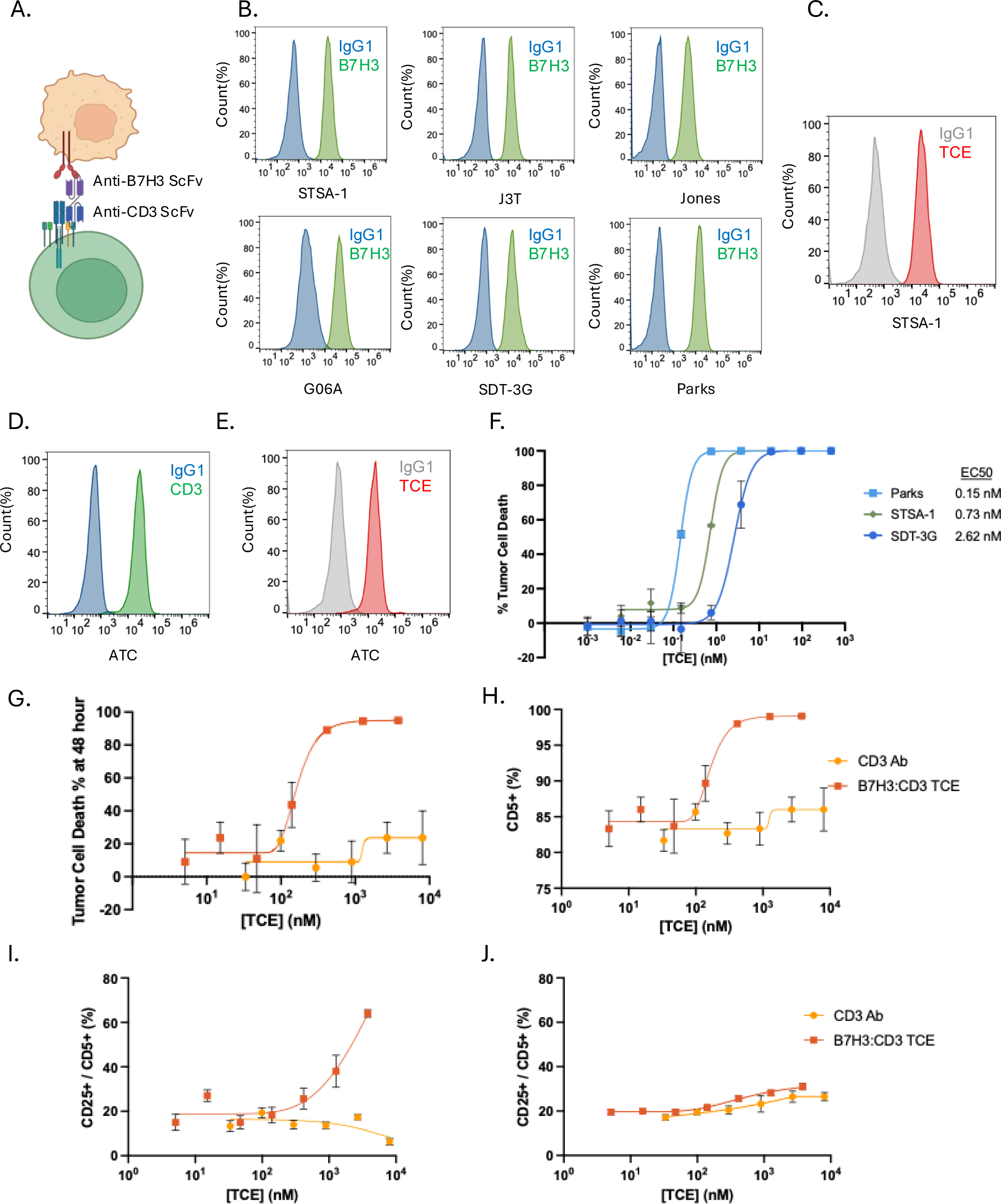
Molecular Characterization and Antigen-Dependent Activation of B7H3:CD3 Bi-specific T-cell engager (TCE) **(A)** Structural schematic of the canine B7H3:CD3 TCE **(B)** B7H3 target antigen expression across a panel of aggressive canine malignancies, including soft tissue sarcoma (STSA-1), glioblastoma (G06A, SDT-3G, J3T), and oral melanoma (Jones and Parks cell lines). **(C)** TCE binding on STSA-1 canine sarcoma cells. **(D–E)** Effector cell engagement: **(D)** Confirmation of CD3 expression on activated canine T cells (ATCs). **(E)** TCE binding on ATCs **(F)** Dose-response curves demonstrating the cytotoxic potential of the TCE across diverse canine malignancies, **(G-H)** Flow cytometry-based cytotoxicity assay at Day 2 **(G)** Dose-response curve **(H)** CD5^+^ T cell events **(I-J)** CD25 expression on T cells. **(I)** In co-culture of STSA-1 and ATC with TCE. **(J)** In ATC only with TCE (absence of target cells).

To validate the therapeutic potential of the B7H3:CD3 TCE, we then characterized its performance across a panel of canine malignancy cell lines. First, we confirmed high B7H3 expression in canine STS, glioblastoma (GBM), and oral melanoma (Fig. 1B). The purity of the TCE was validated by SDS-PAGE and absolute size-exclusion chromatography (Fig. S1). The B7H3:CD3 TCE demonstrated dual-affinity for the B7H3 antigen on STS and for the canine CD3 receptor on activated T cells (Fig. 1C-E), the mechanical prerequisite for the immunological synapse. In T-cell killing (TCK) assays, the TCE demonstrated dose-dependent cytotoxicity across STS, melanoma, and GBM cell lines, with EC50s of 0.72, 0.15, and 2.62 nM, respectively (Fig. 1F-G). While tumor cell lysis begins within 12 hours, maximal cytotoxic effect was observed by 36 hours and sustained up to 120 hours in STS cells (Fig. S2). Notably, the TCE induced CD25 expression on T cells only when cocultured with B7H3^+^ canine STS cells, demonstrating that T-cell activation was antigen-dependent (Fig. 1H-J).

### Establishment of a Multi-Needle Platform for Spatial Pharmacodynamic Mapping

A primary objective of this feasibility study was to determine if an IT injection of B7H3:CD3 TCE at a single dose (7.83 μg /148.2 pmol) could elicit a PD response without triggering the systemic toxicities commonly associated with TCEs. To measure the PD using an internal control and to ensure the geographic precision of our IT delivery, we developed and validated a custom-engineered needle assembly designed to maintain parallel injection tracks triangulated consistently at 1.5 cm apart. This internal geographic control model allowed simultaneous injection of TCE and saline controls into adjacent yet discrete regions of the same tumor, isolating local effects from the background immune landscape of the tumor (Fig. 2).

**Figure 2.**
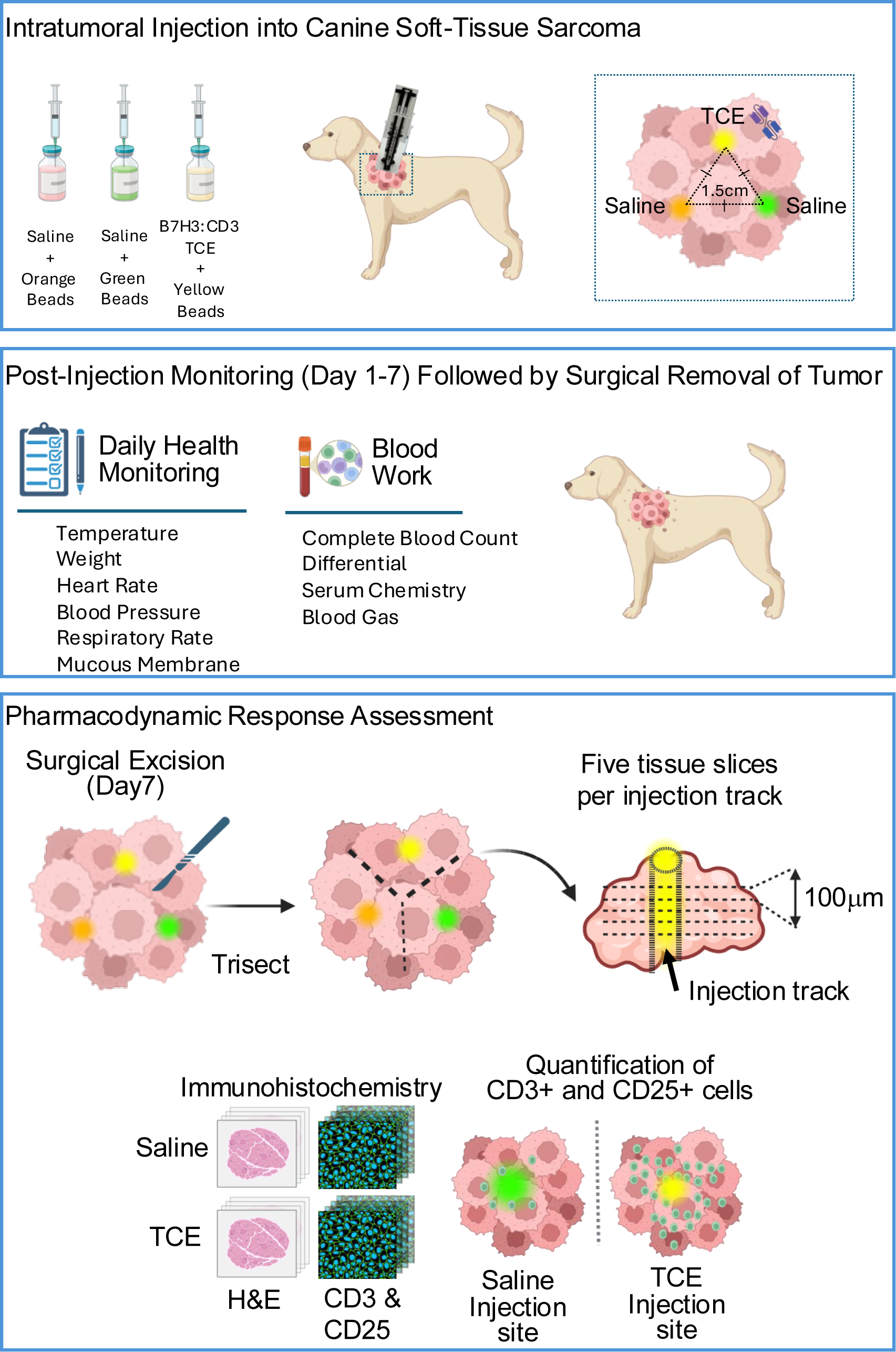
Graphical Abstract of a Canine Clinical Trial of an Intratumoral B7H3:CD3 Bispecific T-cell Engager (TCE). A multi-needle manifold was used for the simultaneous injection of a B7H3:CD3 TCE (comprising a single-chain fragment variable of B7H3 and CD3 linked together) mixed with fluorescent beads, alongside two saline controls, directly into canine soft tissue sarcomas. Following injection, canine patients were monitored for 7 days to assess safety and tolerability. Tumors were then surgically harvested, and H&E staining, as well as immunohistochemistry (IHC) for CD3 and CD25, was performed to evaluate the pharmacodynamic response to the B7H3:CD3 TCE.

The assembly’s spatial accuracy was first confirmed in an *ex vivo* tissue-mimic model using fluorescent micro-beads (Fig. 3 A-B). The beads were spatially defined, indicating that the assembly prevents overlapping needles while maintaining a precise 1.5 cm offset (Fig. 3 C). The syringe holder performed a volumetric column delivery—by withdrawing the needle rather than depressing the plunger during the injection—to distribute the TCE in a 3D vertical column rather than a bolus. This technique ensures that the PD response can be evaluated along the vertical depth of the injection track.

**Figure 3.**
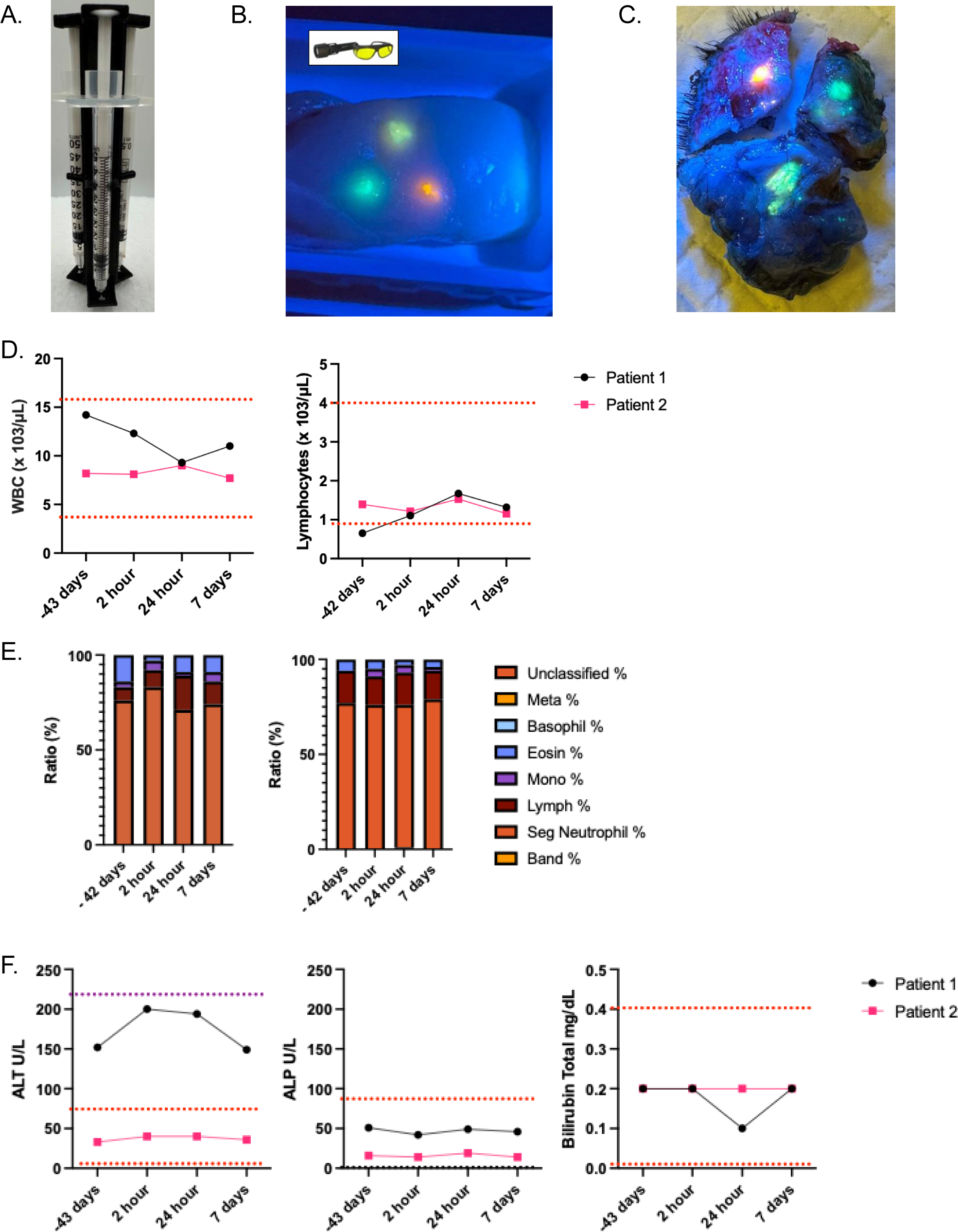
Validation of the Multi-Needle Manifold for Volumetric Intratumoral Delivery. **(A)** The custom-engineered, 3D-printed multi-needle assembly designed for 1-mL insulin syringes to deliver 100 μL volumes at a fixed 1.5cm center-to-center offset. **(B)** *Ex vivo* validation / visualization of three distinct fluorescently labeled microbeads injection in a tissue-mimic model and visualization under blue light **(C)** Clinical application in a canine soft-tissue sarcoma. Representative image of the tumor mass at Day 7 post-injection illustrating the “internal geographic control” model: Needle 1 (B7H3:CD3 TCE), Needle 2 and 3 (Saline controls) **(D)** Longitudinal analysis of complete blood counts (white blood cells and leukocytes) **(E)** Longitudinal analysis of white blood cell differentials **(F)** Longitudinal analysis of serum chemistry and blood gas analysis

### IT Delivery of B7H3:CD3 TCE Demonstrates Tolerability

Using a validated multi-needle platform, the PD and safety of the B7H3:CD3 TCE IT administration was evaluated in two canine STS patients. The patients received diphenhydramine one hour prior to the procedure, followed by a single IT injection of the TCE and saline controls. The patients were monitored, and blood work was collected at the veterinary hospital for the first 24 hours, with subsequent home monitoring by the owners until day 7.

Longitudinal clinical monitoring of the canine subjects confirmed tolerability. In both canine subjects, no adverse events above VCOG-CTCAE Grade 1 were observed throughout the 7-day observation period (19). Vital signs, including body temperature, heart rate, and respiratory rate, remained within normal physiological ranges, with no signs of pyrexia, hypotension, or tachycardia—common clinical signs of TCE systemic toxicity—suggesting that TCE response at this dose was functionally confined to the TME (Fig. S3). Systemic safety was further validated through evaluation of complete blood counts, white blood cell differentials, and serum chemistry. No evidence of systemic leukopenia, neutropenia, or lymphopenia or hematic distress (ALT, ALP, or bilirubin) was observed following administration of the TCE (Fig. 3 D-F, Table S1-4). This indicates that, at this dose, the localized delivery of the B7H3:CD3 TCE did not induce systemic metabolic distress or on-target/off-tumor toxicity.

### B7H3:CD3 TCE Induces a Spatially Limited T-cell Accumulation at Day 7

To assess the PD response, the tumors were harvested 7 days post-injection and trisected to isolate each injection track. To ensure sufficient sampling for statistical analysis, five slices were collected at 100 µm intervals. IHC staining for CD3 and CD25 was performed to evaluate the T-cell response, with CD3^+^ and CD25^+^ cell populations quantified using MATLAB’s Imaging Processing Toolbox (Fig. 2).

On day 7 post-injection, while H&E staining showed no gross architectural changes (Fig. 4A), quantitative analysis of CD3^+^ IHC staining within a 500 μm radius of the injection track demonstrated that the TCE injection sites had a significant five-fold increase in CD3^+^ T-cell density compared to saline controls (**p<0.01 for patient 1, *p<0.05 for patient 2; Fig. 4B-D). Despite local T-cell enrichment, there was no statistical difference in the absolute number of activation markers (CD25^+^) or in the ratio of activated cells (CD25^+^/CD3^+^) between TCE and saline control tracks (Fig S4). Given the transient nature of CD25 expression and the constraint of assessing only one surgical experimental timepoint, the biologic significance of the CD25 staining will require future studies. Nevertheless, the increased T-cell abundance confirms an IT TCE-induced T cell response.

**Figure 4.**
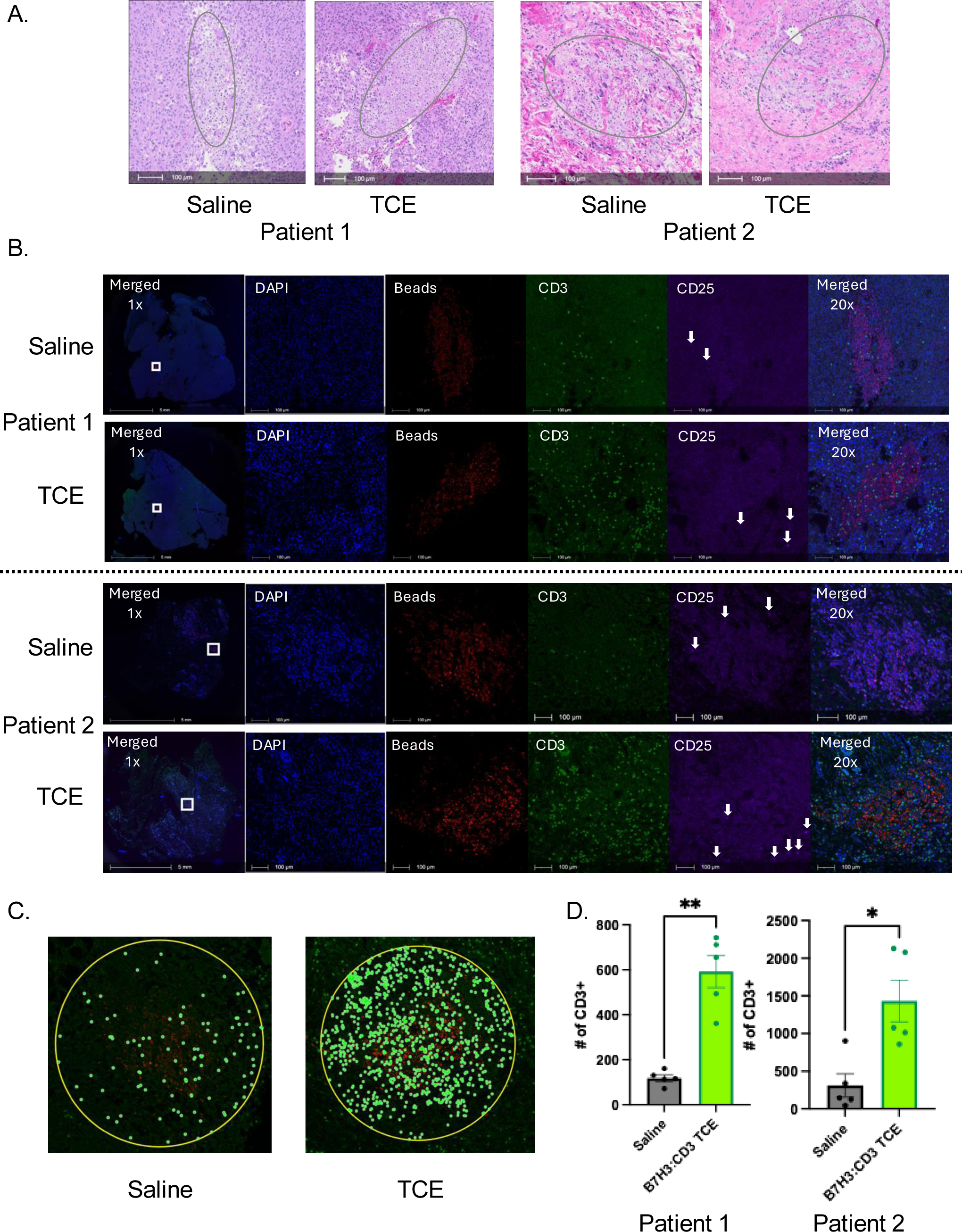
Spatial Pharmacodynamics and Quantitative Mapping of T-cell Sequestration. **(A)** Representative images of H&E centered on the injection sites (grey circle indicates the area of fluorescent beads, which show empty dots in H&E image) **(B)** Representative multi-channel images centered on the injection tracks (identified by fluorescent micro-beads). White square shows the region of the magnified area of interest. White arrows point CD25^+^ cells. **(C)** High-resolution images showing the standardized quantification methodology. Yellow circles define the 500-μm regions of interest used for automated cell counting, while green circles identify the individual CD3^+^ cells. **(D)** Total CD3^+^ T-cell counts in two canine STS patients. **p<0.01, *p<0.05

## Discussion

This feasibility study demonstrates that IT injection of a B7H3:CD3 TCE at a dose of 7.83 μg (148.2 pmol) induces a five-fold increase in CD3^+^ T-cell density at the injection site at day 7 (Fig 4D). The robust accumulation aligns with our *in vitro* data, which showed T-cell expansion and activation at day 2 (Fig. 1H & I). The transition to an *in vivo* environment necessitates a longer observational window to account for the time required for effector cell recirculation, extravasation, and activation within a complex tissue architecture. We selected day 7 as the primary endpoint to capture these spatial-temporal dynamics.

A key finding was the lack of CD25^+^ expression at the day 7 snapshot, despite robust T-cell accumulation. This phenotype presents several mechanistic possibilities. First, given the transient nature of CD25^+^ expression, a peak of activation may have occurred earlier and subsided by the day 7 snapshot (20). Second, the canine STS immunosuppressive TME may impose a metabolic or chemical brake that prevents full T-cell activation (21). Finally, while signal 1 (TCR engagement via the B7H3:CD3 bridge) is sufficient to drive T-cell localization, the single IT dose may be insufficient to sustain effector function without exogenous co-stimulation.

The other notable finding was the lack of T-cell migration across the 1.5 cm gap between injection tracks. By day 7, CD3^+^ cell accumulation remained within the TCE-treated zone, with low CD3^+^ cells in adjacent saline control regions. This phenomenon may be driven by more than one mechanism. First, the dense, extracellular matrix-enriched stroma of the sarcoma may act as a physical cage, which limits T-cell motility from—and creates an immune-sink phenotype at—the TCE injection site (22). Alternatively, this desmoplastic stroma may render the neighboring saline injection regions actively immune-excluded, which harbors biophysical or biochemical barriers that bar T-cell entry, regardless of the activation state (14, 22). Finally, the absence of a broader inflammatory cytokine gradient or chemokine signaling may deprive the recruited T cells of the directional cues necessary to exit the site of initial engagement and penetrate “cold” tumor regions (23, 24).

The localized PD response is ideal for limiting systemic toxicity and provides a roadmap for enhancing the therapeutic index. Based on these results, we have initiated a multi-center canine clinical trial through the Companion Animal Studies for Translational Research Alliance. This follow-up study seeks to improve efficacy while maintaining safety through two primary strategies: 1) co-injection of a TCE that complements CD3 activation with CD28 co-stimulation to promote the T-cell cytoskeletal remodeling and cytokine secretion (e.g., IL-2, IFN-γ) (25); 2) optimization of the number of injections and injection spacing to maximize the tumor volume exposed to the therapeutics.

While IT-injected TCEs are not intended to replace surgery, we envision IT immunotherapy as an adjuvant treatment injected into peri-tumoral tissue following gross total or partial resection to reduce the risk of local recurrence. Neo-adjuvant IT TCEs could also benefit patients with tumors in locations that preclude gross total resection or reduce the risk of surgical morbidity or provide palliation through mass reduction. In all scenarios, optimizing dose and injection spacing in large animals with autochthonous tumors and intact immune systems will increase the likelihood of success in human clinical trials.

In summary, this study establishes that IT delivery of B7H3:CD3 TCEs facilitates local T-cell accumulation without inducing systemic cytokine release or organ toxicity. However, the localized nature of this response suggests that anti-tumor efficacy in bulky solid tumors may require multiple needle injections to ensure volumetric coverage and combinatorial strategies—such as co-stimulation—to unlock the full potential of localized T-cell engagement within the constraints of the TME.

## Materials and Methods

### Bispecific T-cell Engager Production and Purification

The B7H3:CD3 TCE was produced via lentiviral transduction of 293-F cells. Cell culture supernatants were harvested, and the TCE was purified using immobilized metal affinity chromatography followed by size-exclusion chromatography (SEC). Final product quality was validated for purity and endotoxin levels (<0.1 EU/mL) to ensure suitability for *in vivo* administration.

### TCE Stability Test

Protein stability was assessed via analytical SEC. Chromatographic profiles of the TCE stored at 4°C for 5 weeks were compared against freshly thawed aliquots to ensure structural integrity.

### Cell phenotyping

Target cell phenotyping was performed by staining canine cell lines for B7H3 expression (BioLegend, clone MIH42).

### TCE Binding Analysis

Canine tumor cells and activated T cells (ATCs) were incubated with the TCE; binding was detected using an anti-His-AF647 secondary antibody (GenScript) targeting the C-terminal His-tag and analyzed via NovoCyte flow cytometry.

### T-cell activation assay in T-cell killing assay

iRFP-labeled STSA-1 cells were co-cultured with canine ATCs and a five-fold dose titration of the TCE. Tumor cell viability (iRFP signal) was monitored longitudinally using an Incucyte SX5 (Sartorius AG). T-cell activation was quantified by CD25 (eBiosciences) expression via flow cytometry (NovoCyte) in the presence and absence of B7H3^+^ target cells to confirm antigen-dependent engagement.

### Design and Fabrication of the Multi-Needle Manifold

The injection assembly was designed to accommodate three 1-mL insulin syringes in a fixed linear array with a 1.5 cm center-to-center distance between needle tracks. This geometry was optimized to ensure that the individual 100 μL injection volumes remained spatially distinct within the tumor.

### Volumetric Columnar Injection Technique

To ensure a 3D vertical “track” of injection, which allows for PD response evaluation across the full vertical axis in the tumor, the operator held the assembly and steadily withdrew the plungers to deliver a total volume of 100 μL per needle. Following administration, canine subjects were monitored in an intensive care unit for up to 24 hours for acute adverse events. Longitudinal safety was assessed via a combination of daily clinical observations (initially by veterinary hospital staff and subsequently by the patient’s owners), complete blood counts, and serum chemistry panels performed at scheduled intervals post-injection.

### Canine Tissue Processing and H&E

Seven days post-injection, tumor tissues were harvested, formalin-fixed, and processed to paraffin on an automated tissue processor. Infiltrated tissues were embedded in paraffin, sectioned at 4-5 um onto charged slides, baked at 60°C, stained with routine H&E, and coverslipped with permanent mounting media. Whole slide digital imaging was performed on an Olympus VS200 at 20X.

## Acknowledgements

The authors would like to thank the canine patients and their owners for enrolling in this clinical trial; Jason Leubner for critical administration support; Surojit Sarkar and Vandana Kalia for reviewing the manuscript; the Experimental Histopathology Shared Resource, RRID:SCR_022612, of the Fred Hutchinson Cancer Center/University of Washington/Seattle Children’s Cancer Consortium (P30 CA015704), particularly Amanda Koehne, Stephanie Weaver, and Paul Kong for their continued support on H&E and IHC; Connor Burns at Presage Bioscience for the initial brainstorming of IHC strategy; Dawn Duval, Dow Steven, and Rupa Idate at Colorado State University for canine STS and oral melanoma cell lines; and Peter Dickinson and Daniel York at University of California-Davis for canine glioblastoma cell lines.

## Contributors

Conceptualization: YS, JMO. Investigation: YS, LSA, KB, SMM, EJG, AMW, SC, IB, NMH, JH, JF. Visualization: YS, SMM, AMW, EJG. Funding Acquisition: YS, JMO. Project administration: YS. Writing (original draft): YS. Writing (review/editing): YS, LSA, KB, SMM, EJG, AMW, SC, IB, NMH, JH, CD, PM, JPP, JF, JMO.

## Data availability statement

All data associated with this study are present in the paper or the supplementary materials. Request for additional information, materials, and/or script related to this study should be addressed to the corresponding author and may be fulfilled through a material transfer agreement.

## Ethics statements

### Animal Welfare and Ethical Approval

All clinical procedures were performed in accordance with the Washington State University (WSU) Institutional Animal Care and Use Committee (IACUC) guidelines under approved protocol 7106. This study involved client-owned, spontaneous canine cancer patients. All diagnostic and therapeutic interventions were conducted at the WSU College of Veterinary Medicine under the direct supervision of board-certified veterinary oncologists and small animal surgeons.

### Informed Owner Consent

Prior to enrollment, the owners of the canine subjects were provided with a detailed informed consent form outlining the study’s objectives, potential risks, and the experimental nature of the B7H3:CD3 TCE. Participation was entirely voluntary, and owners retained the right to withdraw their animals from the study at any time. All patients received standard supportive care throughout the duration of the trial.

---

**Supplementary Figure 1:**
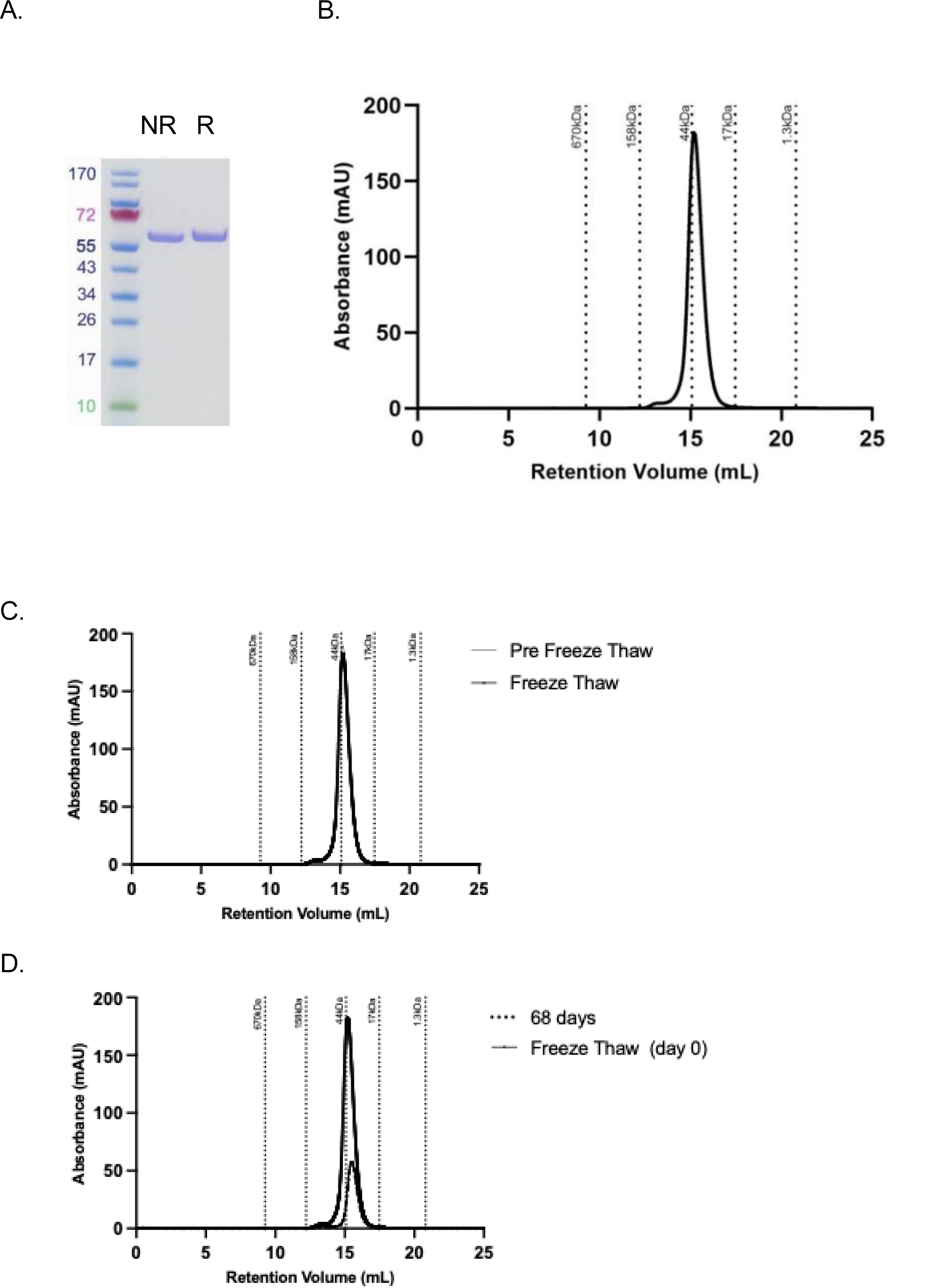
**(A)** SDS-PAGE of Protein. Left lane: non-reduced (NR). Right lane: Reduced (R) **(B)** Absolute Size Exclusion Chromatography (aSEC) of Protein. **C)** aSEC profile overlay of TCE before/after freeze and thaw. **(D)** Overlay of aSEC profile of TCE between day 0 and day 68. Note that day 68 day used 150 μL sample whereas day 0 used 500μl.

**Supplementary Figure 2:**
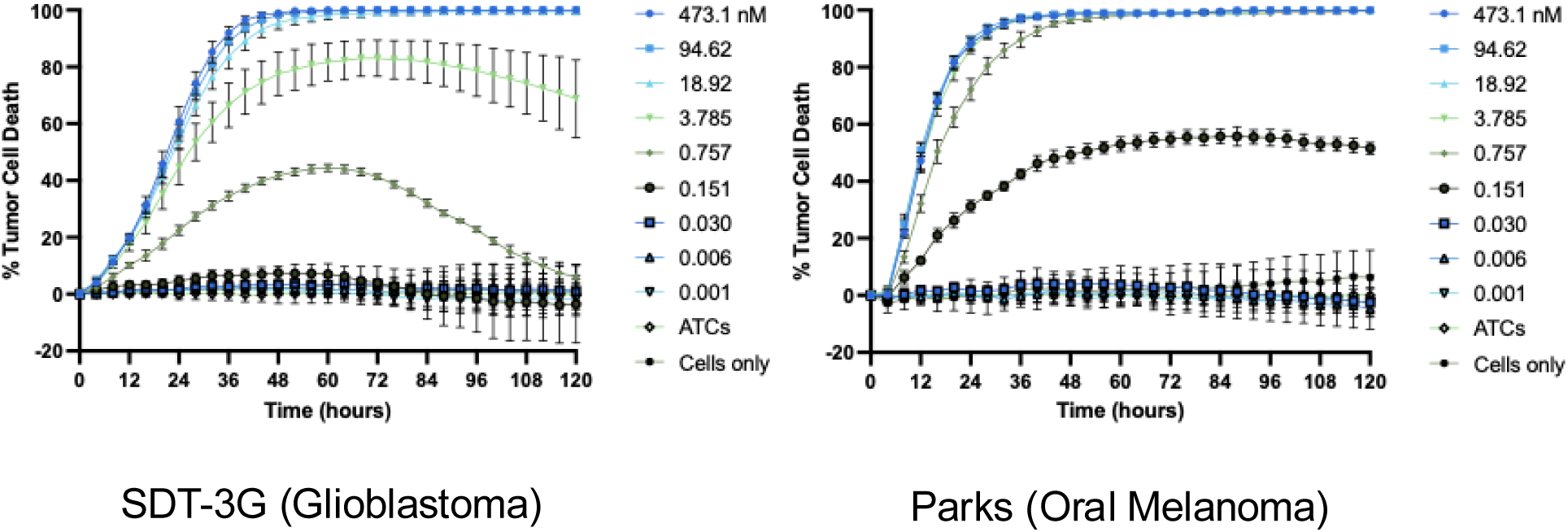
Real-time kinetics of redirected T-cell lysis: Continuous monitoring of SDT-3G and Parks cell death via Incucyte live-cell imaging over 120 hours.

**Supplementary Figure 3:**
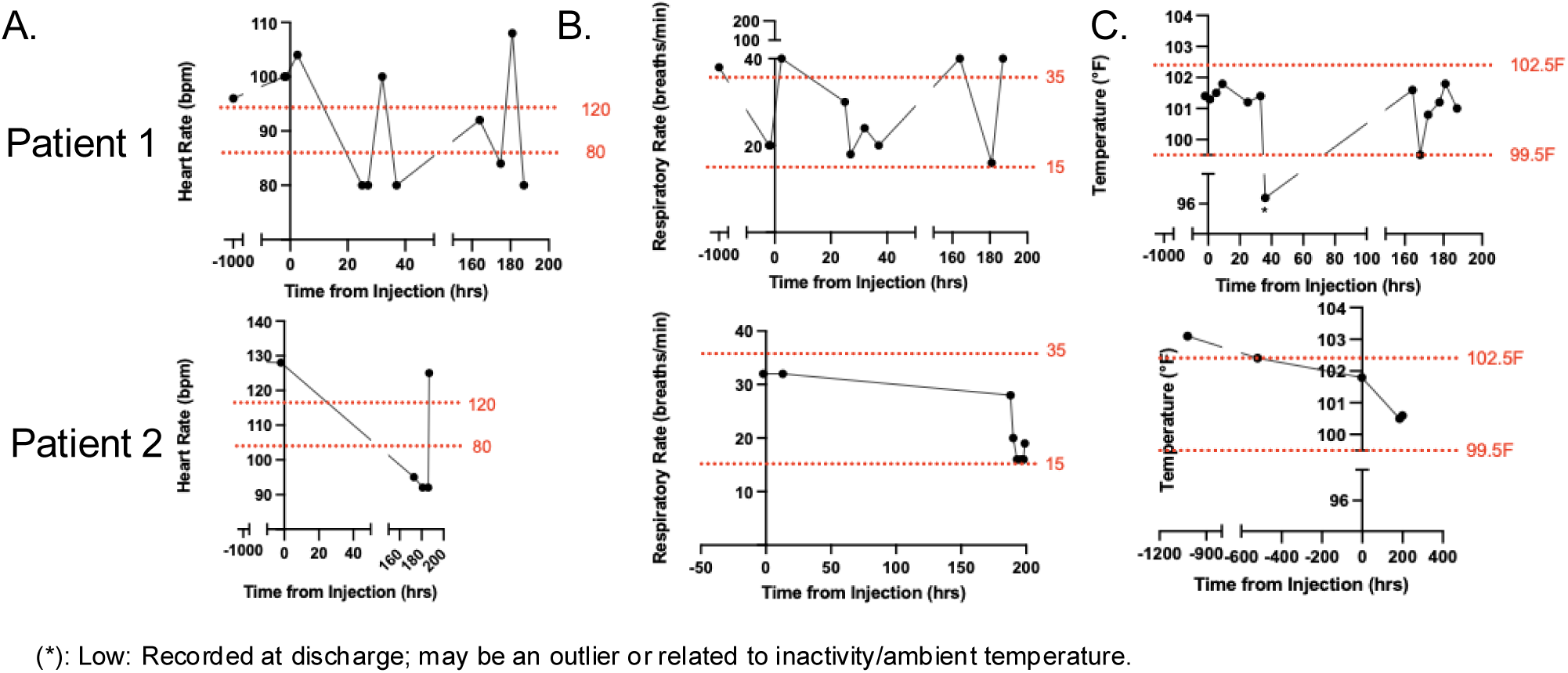
Longitudinal measurements of heart rate, respiratory rate, and body temperature. Data represent observations collected over 7 days with the pre-injection health screening visit as a baseline to monitor physiological trends. Red lines show reference range.

**Supplementary Figure 4:**
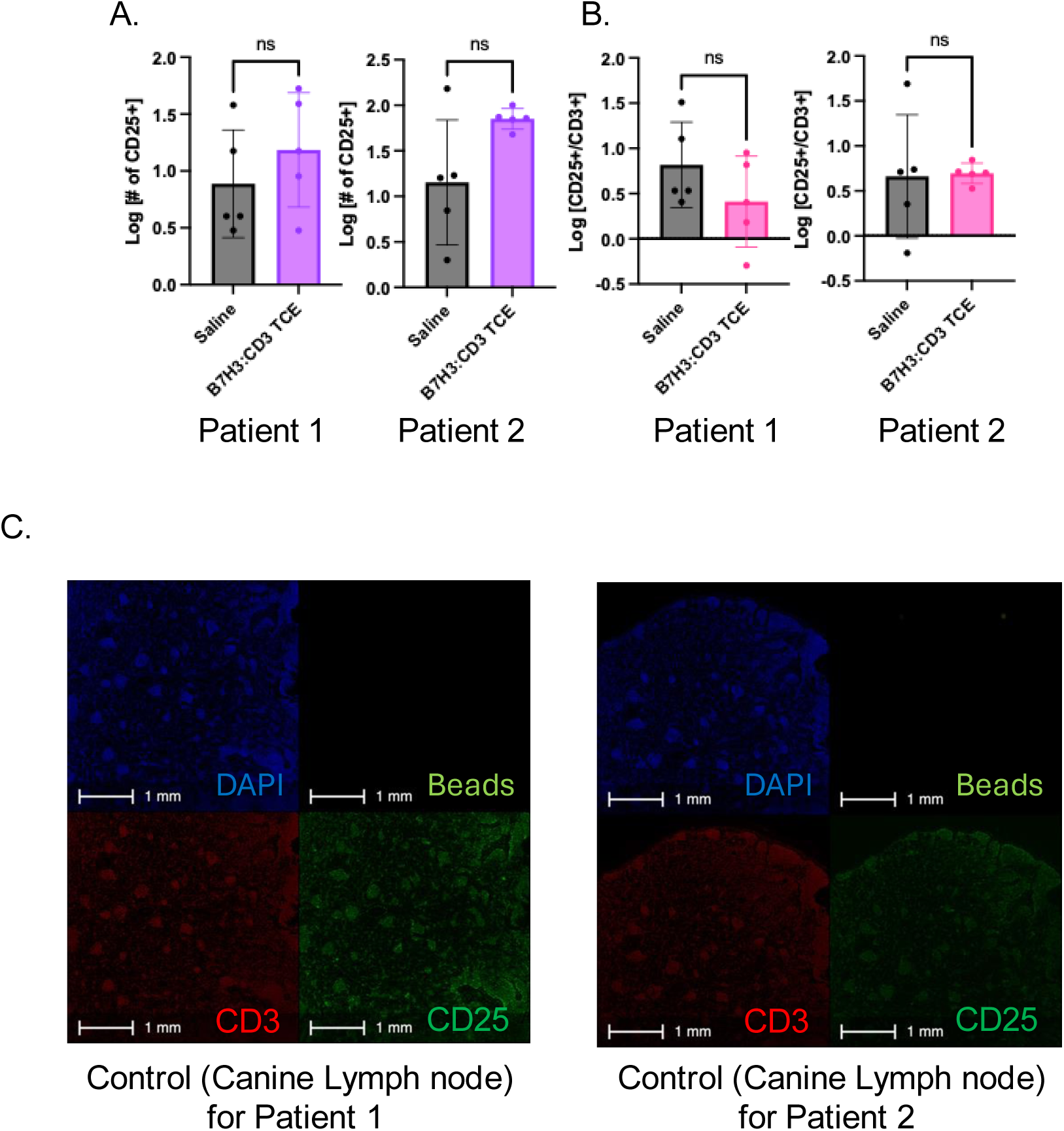
Quantification of CD25^+^ staining on canine STS patients with Validation of Immunohistochemical Staining Protocols in Canine Lymph Node Positive Controls. **A.** Total CD25^+^ T-cell counts in two canine STS patients. **B.** CD25^+^/CD3^+^ ratio in two canine STS patients. **C.** Representative IHC images of reactive canine lymph nodes used as positive controls for CD3 and CD25 antibody titration and validation. Patient 1

**Supplementary Table 1:** Longitudinal Hematologic Stability and Complete Blood Count (CBC) Works for a canine STS patient 1 following localized B7H3:CD3 TCE administration.

| Test | Reference Interval | -42 days | 2 hour | 24 hour | 7 days |
| --- | --- | --- | --- | --- | --- |
| WBC (x 10<sup>3</sup>/μL) | 4.5 - 16 | 9.3 | 12.3 | 9.3 | 11 |
| Seg# (x 10<sup>3</sup>/μL) | 2.8 - 13.4 | 7.068 | 10.209 | 6.603 | 8.14 |
| Band# (x 10<sup>3</sup>/μL) | 0 - 0.3 | 0 | 0 | 0 | 0 |
| Lymph# (x 10<sup>3</sup>/μL) | 0.9 - 4.0 | 0.651 | 1.107 | 1.674 | 1.32 |
| Mono# (x 10<sup>3</sup>/μL) | 0 - 1.3 | 0.279 | 0.615 | 0.186 | 0.55 |
| Eosin# (x 10<sup>3</sup>/μL) | 0 - 1.2 | 1.302 | 0.369 | 0.837 | 0.99 |
| RBC (x 10<sup>6</sup>/μL) | 5.2 - 8.4 | 7.48 | 6.58 | 6.84 | 7.67 |
| Hgb (g/dL) | 12 - 20 | 17 | 15 | 16.1 | 17.4 |
| PCV (%) | 36 - 56 | 51 | 42 | 45 | 52 |
| Platelets (x 10<sup>3</sup>/μL) | 160 - 500 | 294 | 213 | 236 | 236 |

**Supplementary Table 2:** Longitudinal Hematologic Stability and Complete Blood Count (CBC) Works for a canine STS patient 2 following localized B7H3:CD3 TCE administration.

| Test | Reference Interval | -43 days | 2 hour | 24 hour | 7 days |
| --- | --- | --- | --- | --- | --- |
| WBC (x 10<sup>3</sup>/μL) | 4.5 - 16 | 8.2 | 8.1 | 9 | 7.7 |
| Seg# (x 10<sup>3</sup>/μL) | 2.8 - 13.4 | 6.314 | 6.156 | 6.75 | 6.083 |
| Band# (x 10<sup>3</sup>/μL) | 0 - 0.3 | 0 | 0 | 0.09 | 0 |
| Lymph# (x 10<sup>3</sup>/μL) | 0.9 - 4.0 | 1.394 | 1.215 | 1.53 | 1.155 |
| Mono# (x 10<sup>3</sup>/μL) | 0 - 1.3 | 0 | 0.324 | 0.36 | 0.154 |
| Eosin# (x 10<sup>3</sup>/μL) | 0 - 1.2 | 0.492 | 0.405 | 0.27 | 0.308 |
| RBC (x 10<sup>6</sup>/μL) | 5.2 - 8.4 | 8.02 | 7.53 | 5.76 | 7.18 |
| Hgb (g/dL) | 12 - 20 | 19.7 | 18.4 | 14.6 | 18.1 |
| PCV (%) | 36 - 56 | 59 | 53 | 42 | 52 |
| Platelets (x 10<sup>3</sup>/μL) | 160 - 500 | 222 | 184 | 172 | 193 |

**Supplementary Table 3:** Comprehensive biochemical profiling of a canine STS subject at baseline (Day -42), and 2-hour, 24-hour, and 7-day intervals following intratumoral B7H3:CD3 TCE injection for Patient 1.

| Test | -42 days | 2 hour | 24 hour | 7 days | Reference Intervals |
| --- | --- | --- | --- | --- | --- |
| ALT U/L | 152 | 200 | 194 | 149 | 8 - 77 |
| ALP U/L | 51 | 42 | 49 | 46 | 0-87 |
| Cholesterol mg/dL | 208 | 222 | 241 | 246 | 131-346 |
| BUN mg/dL | 15 | 19 | 16 | 12 | 10 - 26 |
| Creatinine mg/dL | 1.2 | 1.2 | 1 | 1 | 0.64-1.46 |
| Glucose mg/dL | 101 | 99 | 57 | 92 | 75-120 |
| Total Protein g/dL | 6.3 | 5.7 | 6.1 | 6.4 | 5.4-7.1 |
| Albumin g/dL | 3 | 2.8 | 3 | 3 | 2.8-3.8 |
| Globulin g/dL | 3.3 | 2.9 | 3.1 | 3.4 | 2.2-3.6 |
| Calcium mg/dL | 10 | 10 | 10 | 10.3 | 9.2-10.9 |
| Phosphorus mg/dL | 2.8 | 2.9 | 4.3 | 3.6 | 2.2-5.9 |
| Bilirubin Total mg/dL | 0.2 | 0.2 | 0.1 | 0.2 | 0-0.4 |
| Sodium mEq/L | 149.1 | 148.8 | 149.6 | 150.8 | 143.1-152.9 |
| Potassium mEq/L | 4.7 | 4.7 | 4.7 | 4.6 | 3.9-5 |
| Chloride mEq/L | 113 | 116 | 114 | 112 | 110-120.5 |
| CO2 mEq/L | 25.2 | 24.6 | 26.2 | 26.1 | 14.4-23.6 |
| Anion Gap | 16 | 13 | 14 | 17 | 13-24 |

**Supplementary Table 4:** Comprehensive biochemical profiling of a canine STS subject at baseline (Day -43), and 2-hour, 24-hour, and 7-day intervals following intratumoral B7H3:CD3 TCE injection for Patient 2.

| Test | -43 days | 2 hour | 24 hour | 7 days | Reference Intervals |
| --- | --- | --- | --- | --- | --- |
| ALT U/L | 33 | 40 | 40 | 36 | 8 - 77 |
| ALP U/L | 16 | 14 | 19 | 14 | 0-87 |
| Cholesterol mg/dL | 215 | 240 | 236 | 231 | 131-346 |
| BUN mg/dL | 24 | 14 | 15 | 16 | 10 - 26 |
| Creatinine mg/dL | 1.2 | 0.9 | 0.9 | 1.1 | 0.64-1.46 |
| Glucose mg/dL | 102 | 101 | 104 | 109 | 75-120 |
| Total Protein g/dL | 6 | 5.8 | 5.8 | 5.8 | 5.4-7.1 |
| Albumin g/dL | 3.1 | 3 | 3 | 3 | 2.8-3.8 |
| Globulin g/dL | 2.9 | 2.8 | 2.8 | 2.8 | 2.2-3.6 |
| Calcium mg/dL | 10.2 | 10.1 | 10 | 9.7 | 9.2-10.9 |
| Phosphorus mg/dL | 3.8 | 4.8 | 4.4 | 3.2 | 2.2-5.9 |
| Bilirubin Total mg/dL | 0.2 | 0.2 | 0.2 | 0.2 | 0-0.4 |
| Sodium mEq/L | 147.1 | 147.2 | 146.9 | 145.8 | 143.1-152.9 |
| Potassium mEq/L | 4.3 | 4.6 | 4.4 | 4.3 | 3.9-5 |
| Chloride mEq/L | 115 | 114 | 115 | 116 | 110-120.5 |
| CO2 mEq/L | 21.3 | 23.3 | 23.9 | 18.4 | 14.4-23.6 |
| Anion Gap | 15 | 15 | 12 | 16 | 13-24 |

