## Supplementary Methods for "Intratumoral B7H3:CD3 Bispecific T-cell Engager Drives Localized T-cell Accumulation in Canine Sarcoma Patients"

**Cell culture -** Canine soft-tissue tumor cell line (STSA-1) was cultured in DMEM with 15% FBS, Glutamax, MEM Vitamin, NEAA, and sodium pyruvate in 5% CO2-containing incubators. Canine Glioblastoma cell lines (G06A, SDT3G and J3T) and oral melanoma cell lines (Jones and Park) were cultured in DMEM with 10% FBS and Glutamax in 5% CO2-containing incubators.

**Bi-specific T-cell Engager Design, Production, and Purification -** Cells (Freestyle 293-T, ATCC #CRL3216) were transfected with DNA and packaging vectors to make lentiviruses encoding the protein of interest using the Daedalus System, as previously described by Bandaranayake et al. (Nucleic Acids Research, 2011).[1] Freestyle 293-F cells (Invitrogen, #R79007) were transduced with lentivirus harvested from the 293-T cells encoding the protein of interest. Cells were grown aseptically for two weeks in 293-F expression medium (Gibco # 12338018) until they reached a viability of 80% and a cell density of $3.5e^6$ cells/mL. The culture was centrifuged at 3,470 rcf for 20 minutes in a Sorvall Lynx 6000 centrifuge and the supernatant, containing the protein, was collected.

The protein was purified via immobilized metal affinity chromatography (IMAC) and size exclusion chromatography (SEC) on an Äkta Pure 25 fast protein liquid chromatography (FPLC) system (Cytiva). The protein was analyzed by UV/Vis (Thermo Scientific, NanoDrop One), dynamic light scattering (Unchained Labs, Stunner), and sodium dodecyl sulfate-polyacrylamide gel electrophoresis (SDS-PAGE). The protein's size and purity was also verified using analytical SEC (aSEC) on a Superdex 200 Increase 10/300 GL column (Cytiva, #28990944). The protein was also tested for endotoxins, and verified to be endotoxin-free, with an Endosafe Nexgen-PTS portable endotoxin tester (Charles River, #PTS150). The protein was then aliquoted, flash-frozen in liquid nitrogen, and stored at -80 $^{o}$C.

**TCE Stability Test -** The protein was tested for stability via aSEC. An aliquot of protein was flash-frozen in liquid nitrogen, allowed to thaw at room temperature, and run on the aSEC column. The pre and post freeze/thaw aSECs were overlayed. There was no difference between the aSEC of the protein before and after freezing, indicating that the protein was stable. After the initial aSEC was run, the protein was stored at 4 $^{o}$C for 5 weeks and another aSEC was run. There was no difference between the aSECs indicating that the protein was stable for at least 5 weeks at 4 $^{o}$C.

**Cell phenotyping -** Canine cells were blocked with EasySep (STEMCELL Technologies) with 3% Normal Rat Serum for 20 minutes at 4 $^{o}$C. Antibodies to cell surface marker were also diluted in the cell blocking buffer. Cells were then stained for 30 minutes at 4 $^{o}$C. Samples were analyzed on the Novocyte 3000 using Novosoft software for analysis. Cells were stained for B7H3 (Biolegend, clone MIH42) expression.

**TCE Binding Analysis -** TCEs were used at 10 μg/mL and stained 100K cells in 50 μL volume. After incubating cells with TCEs, the cells were washed 2xs in Easysep, then incubated with the anti-HIS-AF647 secondary (Genescript) for 30 minutes at 4 $^{o}$C. Cells were then analyzed on the Novocyte 3000 using Novosoft analysis software.

**T-cell activation assay in T-cell killing assay -** Frozen canine Peripheral Blood Molecular Cells (PBMCs) were thawed in RPMI with 10% FBS media supplemented with non-essential amino acids (Gibco), 2-mercaptoethanol (Gibco) and Penicillin/Streptomycin (Gibco), then plated in the same media into flat bottom, 96-well plates with titrated TCEs. PBMCs were seeded at 50K/well while iRFP-labeled STSA-1 were seeded at 10K/well. Combined cells were cultured for 4 days at 37 $^{o}$C, 5% CO2. TCEs were titrated with either PBMCs alone or PBMCs with STSA-1 tumor cells. After culture, cells were washed 2xs in 3 mM EDTA in PBS, then with 50 µL/well trypsin for five minutes. After five minutes, FBS was added at 50 µL/well. Cells were then washed 1x in EasySep, then resuspended in EasySep with 3 % NRS. Cells were stained for CD5 and CD25. After staining, cells were analyzed for iRFP+ tumor cells to determine tumor cell inhibition, CD5, and CD25 to score activation of PBMCs. Cells were analyzed on the Novocyte 3000 using Novosoft software.

**T-cell killing assay -** Fluorescently labeled canine tumor cells were cocultured in triplicate with a five-fold dose titration of TCE and canine ATCs at a 5:1 effector to target ratio. Plates were imaged on an Incucyte SX5 (Sartorius AG) every 4 hours for up to 5 days and analyzed for loss of fluorescent tumor cells. Data was analyzed using GraphPad Prism10 to generate dose response curves.

**Design and Fabrication of the Multi-Needle Manifold –** To achieve standardized volumetric delivery and ensure non-overlapping vertical column distribution, a custom-engineered injection assembly was developed. The assembly was initially modeled in Onshape and subsequently fabricated using a Markforged Mark Two 3D printer with Onyx resin (stl files available upon request). Specifically, the injection assembly was designed to accommodate three 500-µL insulin syringes in a fixed linear array with a 1.5 cm center-to-center distance between needle tracks. This geometry was optimized to ensure that the individual 100 µL injection volumes remained spatially distinct within the tumor.

**Ex-Vivo Validation of Spatial Precision -** Prior to *in vivo* administration, the spatial accuracy of the manifold was validated using a tissue-mimic model (*ex vivo* chicken breast). Three distinct fluorescently labeled medical-degree microbeads (yellow, green, and orange under blue light), CIVO-Glo (Presage Bio Science), were loaded into the manifold syringes. A retrograde "columnar" injection was performed as described below. The tissue was imaged under blue-light, and then cross-sectioned to confirm the absence of overlap between injection sites and to verify column injection.

**Volumetric Columnar Injection Technique -** For each canine subject, the tumor site was prepared using standard aseptic techniques. Under clinical observation, the multi-needle array was inserted to the maximal depth of the target tumor mass. To achieve a continuous column injection rather than a localized spherical bolus, a retrograde delivery technique was employed. The operator held the manifold and steadily withdrew the plungers to deliver a total volume of 100mL per needle. This technique created a 3D vertical "track" of B7H3:CD3 TCE or saline to ensure that the pharmacodynamic response could be evaluated across the full vertical axis in the tumor.

**Canine Blood Work -** Prior to injection, and then two hours post injection, blood was pulled from a peripheral vein and submitted for assessment of a complete blood count and serum chemistries. Serum was also frozen back for assessment of cytokines at a later date. A third blood draw was done the following day approximately 24 hours after the first blood draw post injection. A $4^{th}$ blood draw was done 7 days post injection on surgery day.

**Clinical Monitoring -** For each patient a physical exam was done the day of the injection and tumor was measured in 3 dimensions. Baseline blood pressure was recorded and diphenhydramine (1 mg/kg orally) was given 1 hour prior to the injection of the T cell engager intratumorally. The 3-needle injection was given in the center of the tumor and then the patients were monitored for any potential side effects for roughly 18 hours. The first 6-7 hours were close observation and the next 11-12 hours were often in ICU where the patient was monitored overnight. Diphenhydramine administration was repeated at 12 hours and 24 hours post injection. Blood pressure was taken a third time in the morning after the injection (roughly 24 hours post injection) and the mass was measured in three dimensions. At recheck 7 days later when presenting for surgery, the mass was again measured prior to removal and blood pressure was taken.

The patient was monitored constantly by veterinary staff for any signs of developing anaphylaxis. Breathing was visually monitored as well as attitude and appetite to be sure the patient was feeling well. If there were concerns of any sort due to the patient being agitated or less responsive, the heart and lungs were auscultated, and the temperature was taken. The patient was also walked outside if there was concern for development of diarrhea. Patients were sent home with their owners once they had been observed for roughly 24 hours post injection. The owners were advised to monitor any abnormal behaviors in their dog and to contact the veterinarian or technician associated with the trial. They were also instructed to give diphenhydramine if any respiratory or GI signs were noted.

**Canine Tissue Processing and H&E -** Paraffin processing was performed on the Sakura Tissue-Tek VIP with two 1-hour cycles of 100% ethanol dehydration, three cycles of Clear Rite (Thermo Scientific Catalog#: 22046341) and four cycles of paraffin at 60°C (Leica 39601006). Samples were embedded in paraffin on a Tissue Tek TEC embedding station. Paraffin embedded blocks were sectioned on a Leica RM2255 at 4-5 microns, air dried at room temperature overnight and baked at 60°C prior to staining. Slides with paraffin sections were processed using the Sakura Tissue-Tek Prisma automated stainer for Hematoxylin and Eosin (H&E) staining. Slides were deparaffinized and rehydrated in multiple changes of each of the following reagents: xylene (Fisher Scientific, X3P) for 5 minutes, 100% ethanol (Decon Labs, Inc. 2701) for 4 minutes, 95 % ethanol for 4 minutes, water rinse for 1 minute. Slides were stained in hematoxylin (Epredia 72711) for 11 minutes followed by a 1 minute water rinse, a 1 minute clarifier incubation (Epredia 7401), a 2 minute water rinse, Bluing Reagent (Epredia 7301) for 1 minute, a 1 minute water rinse, 95% ethanol for 1 minute followed by Eosin Y Phloxine for 2 minutes. Slides were dehydrated in graded ethanol with two 1 minute 95% ethanol incubation and 3 100% ethanol incubations, and cleared with xylene for at least 4 minutes. Whole slide digital imaging was performed on an Olympus VS200 at 20X.

**Immunohistochemistry -** Formalin-fixed paraffin-embedded tissues were sectioned at 4 microns onto positively charged slides and baked for 1 hour at 60°C. The slides were then dewaxed and stained on a Leica BOND Rx stainer (Leica, Buffalo Grove, IL) using Leica Bond reagents for dewaxing (Dewax Solution), antigen retrieval/antibody stripping (Epitope Retrieval Solution 2), and rinsing after each step (Bond Wash Solution). Antigen retrieval and antibody stripping steps were performed at 100°C with all other steps at ambient temperature.

| Position | Antibody | Clone/ Host | Manufacturer/ Catalog Number | Dilution/ concentration | Secondary | Opal Dye |
|---|---|---|---|---|---|---|
| 1 | **CD3** | CD3-12 / Rat | Serotec / MCA 1477 | 1:250 4ug/ml | ImmPress Rat-HRP | **Opal 690** |

| | | | | | | |
|---|---|---|---|---|---|---|
| 2 | **CD25** | 4C9 / Mouse | Cell Marque / 125M-18 | Pre-dilute 0.2ug/ml | OPAL Ms/Rb-HRP | **Opal 780** |

Endogenous peroxidase was blocked with 3% H2O2 for 5 minutes followed by protein blocking with TCT buffer (0.05M Tris, 0.15M NaCl, 0.25% Casein, 0.1% Tween 20, 0.05% ProClin300 pH 7.6) for 10 minutes. CD3 was applied for 60 minutes followed by Vector ImmPress Goat anti-Rat IgG Polymer Detection secondary antibody application for 20 minutes, and the tertiary TSA-amplification reagent (Akoya OPAL 690 fluor) for 20 minutes. A high stringency wash was performed after the secondary and tertiary applications using high-salt TBST solution (0.05M Tris, 0.3M NaCl, and 0.1% Tween-20, pH 7.2-7.6). The primary and secondary antibodies were stripped with retrieval solution for 20 minutes before repeating the process with the second primary antibody (position 2) starting with a new application of 3% H2O2 for 5 minutes, 10% TCT for 10 minutes, primary antibody for 60 minutes, secondary antibody Akoya Opal anti-Mouse+Rabbit HRP secondary antibody for 20 minutes, Opal TSA-DIG for 10 minutes, followed by the 20 minute stripping step in retrieval solution and application of Opal 780 fluor for 10 minutes with high stringency washes performed after the secondary, TSA DIG, and Opal 780 fluor applications. The stripping step was not performed after the final position. Slides were removed from the stainer and stained with DAPI for 5 minutes, rinsed for 5 minutes, and coverslipped with Prolong Gold Antifade reagent (Invitrogen/Life Technologies, Grand Island, NY). Slides were cured overnight at room temperature, then whole slide images were acquired on the PhenoImager HT 2.0 Imaging System (Akoya Biosciences, Marlborough, MA) using the MOTiF whole slide workflow and the images exported as unmixed QPTIFF images.

### Statistical analysis

In Fig 4 and Supplemental Fig 4, $n = 5$ replicates were used, where each replicate was a slice. Where error bars are shown, all values indicate the means ± SD. Significance was determined using paired t-test in GraphPad Prism.
